# HDX-MS reveals nucleotide-regulated, anti-correlated opening and closure of SecA and SecY channels of the bacterial translocon

**DOI:** 10.1101/595553

**Authors:** Zainab Ahdash, Euan Pyle, William J. Allen, Robin A. Corey, Ian Collinson, Argyris Politis

## Abstract

The bacterial Sec translocon is a multi-component protein complex responsible for translocating diverse proteins across the plasma membrane. For post-translational protein translocation, the Sec-channel – SecYEG – associates with the motor protein SecA to mediate the ATP-dependent transport of unfolded pre-proteins across the membrane. Based on the structure of the machinery, combined with ensemble and single molecule analysis, a diffusional based Brownian ratchet mechanism for protein secretion has been proposed [Allen et al. eLife 2016;5:e15598]. However, the conformational dynamics required to facilitate this mechanism have not yet been fully resolved. Here, we employ hydrogen-deuterium exchange mass spectrometry (HDX-MS) to reveal striking nucleotide-dependent conformational changes in the Sec protein-channel. In addition to the ATP-dependent opening of SecY, reported previously, we observe a counteracting, also ATP-dependent, constriction of SecA around the mature regions of the pre-protein. Thus, ATP binding causes SecY to open and SecA to close, while ATP hydrolysis has the opposite effect. This alternating behaviour could help impose the directionality of the Brownian ratchet for protein transport through the Sec machinery, and possibly in translocation systems elsewhere. The results highlight the power of HDX-MS for interrogating the dynamic mechanisms of diverse membrane proteins; including their interactions with small molecules such as nucleotides (ATPases and GTPases) and inhibitors *(e.g.* antibiotics).

## Main

Protein transport across and into the membranes that surround and sub-divide cells is essential for life. The ubiquitous general secretory (Sec) machinery performs this task in the plasma membrane of bacteria and the endoplasmic reticulum of eukaryotes. A protein-channel through the membrane is formed by the conserved hetero-trimeric core-complex: Sec61αβγ in eukaryotes, and SecYEG in archaea and bacteria^1^. In bacteria, protein secretion is important for cell envelope biogenesis and maintenance, as well as for the delivery of adherence and pathogenic effector proteins to the cell surface. During the process of protein secretion, SecYEG engages with the cytosolic motor ATPase SecA^2^, and together they pass pre-proteins with a short N-terminal cleavable signal sequence across the membrane, whilst still in an unfolded conformation^3,4^.

A structure of the complete SecA-SecYEG complex has been determined with an ATP analogue (ADP-BeF_x_; PDB code 3DIN), as well as with a short pre-protein mimic^5,6^ (PDB code 5EUL; modified in **Figure 1a, b**). No structure of the ADP-bound state has been experimentally determined, however molecular dynamics (MD) simulations and Förster Resonance Energy Transfer (FRET) analyses suggest that ATP and ADP respectively favour open and closed forms of the SecY channel^7,8^. This behaviour was incorporated into a ‘Brownian ratchet’ model to describe the mechanism of ATP driven protein translocation, whereby ATP binding and hydrolysis act to bias pre-protein diffusion in an outward direction. This directionality is further augmented by other factors, including the promotion of pre-protein folding on the outside, but not the inside^9^, and by coupling transport to the trans-membrane proton-motive-force (PMF)^10^.

**Figure 1:**
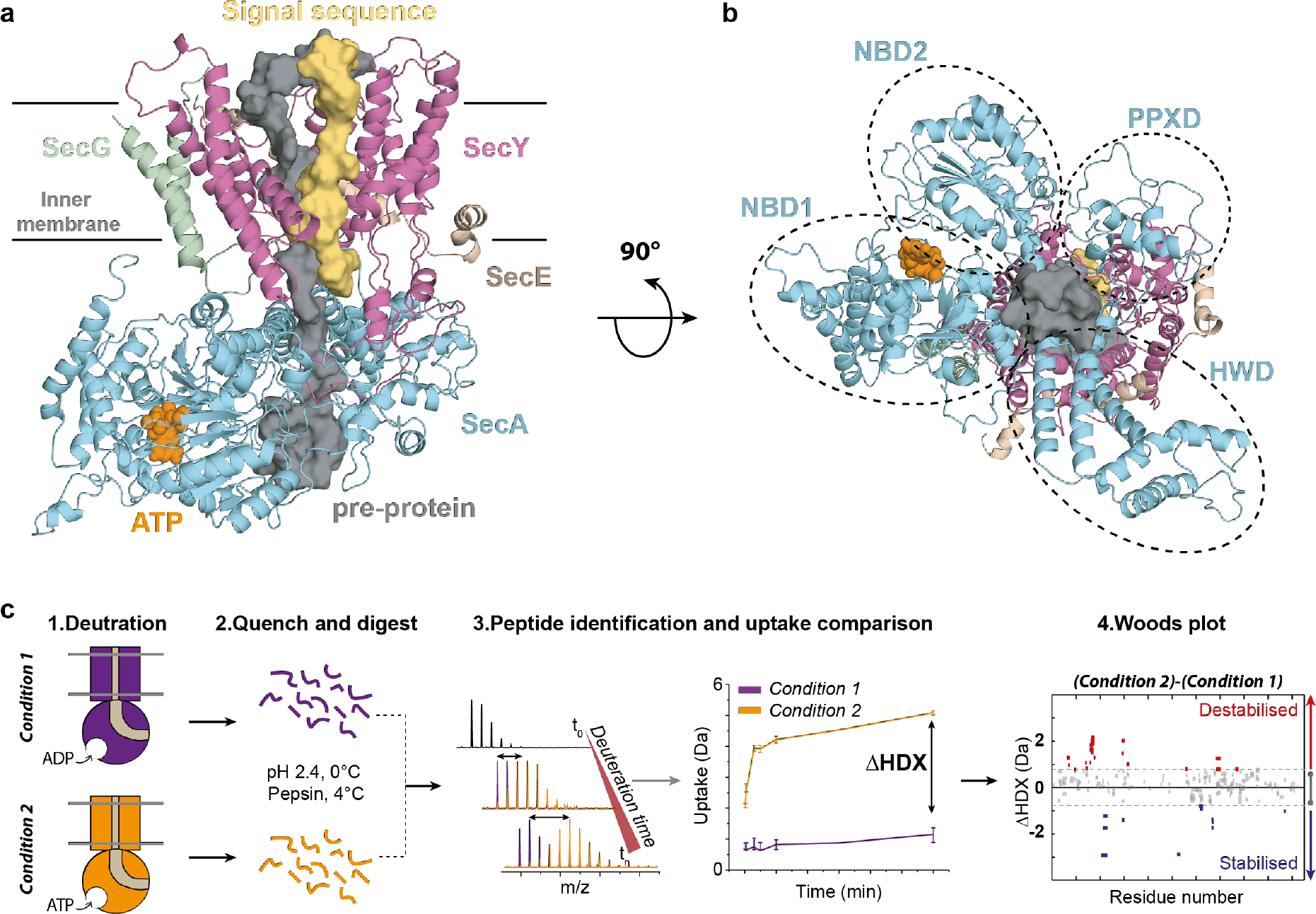
Structure of SecA-SecYEG complex and HDX-MS workflow. (**a**) Structure and subdomains of SecA-SecYEG (based in 3DIN, ^13^with a pre-protein modelled in^26^). The pre-protein, ATP and signal sequence are highlighted in grey, orange and yellow, respectively. (**b**) Top view of the complex highlighting the nucleotide-binding-domains NBD1 and NBD2 as well as PPXD (**P**re-**P**rotein **cross**-linking **D**omain) and HWD (**H**elical **W**ing **D**omain) (**c**) Overview of the HDX process. The sample is prepared in detergents and after addition of nucleotides is incubated in a deuterated solvent (1). Following deuteration of the mixture at different time-points, the HDX reaction is quenched and the protein is digested with pepsin (2). Peptides are separated by liquid chromatography and identified by mass spectrometry (MS). The mass uptake of the protein in different conditions is then compared (3). Peptides with significant difference in deuterium uptake are mapped onto a Woods plot (4).

One apparent contradiction in the Brownian ratchet model is the effect of non-hydrolysable ATP analogues: while they cause the channel through SecY to open-up^8^, they prevent backsliding of trapped translocation intermediates^11,12^. These two observations could be reconciled if SecA itself grips substrates more tightly in its ATP-bound state. SecA forms a clamp for the translocating pre-protein, which could conceivably act in this way (**Figure 1b**)^13^ However, as the this clamp can be fixed in a closed state without preventing transport^14^ the conformational changes involved are likely to be subtle, and not necessarily captured with techniques such as FRET.

Clearly, the key to understanding the mechanism that prevents unfavourable back-sliding and help refine the Brownian ratchet model is to learn more about the dynamic action of SecA-SecYEG throughout the hydrolysis cycle. Here, we exploit recent technical advances in hydrogen-deuterium exchange mass spectrometry (HDX-MS) to probe in exquisite detail the structural differences between the ATP and ADP associated states that underpin function.

HDX-MS has emerged as a non-invasive and highly sensitive method for interrogating the conformational dynamics of membrane proteins and their complexes^15–18^. HDX-MS measures the exchange of backbone amide hydrogen to deuterium at peptide-level resolution^19–21^. The rate of HDX depends on the solvent accessibility, protein flexibility, and hydrogen bonding ^22^. Thus, HDX-MS can monitor the conformational changes of proteins and their complexes by comparing HDX rates between distinct protein states (*e.g.* protein *versus* protein with ligand). Consequently, comparative HDX-MS has emerged as an attractive experimental tool to define conformational changes to complement other, high-resolution techniques. For example, the analysis of the structural consequences of protein-ligand/drug interactions, and nucleotide dependent protein dynamics; ATP/GTP *versus* ADP/GDP^23,24^. Moreover, our group and others have developed and applied HDX-MS to study the conformational diversity of challenging membrane protein systems^15,17,18^.

Here, we use HDX-MS to interrogate nucleotide-induced conformational changes in the multi-component SecA-SecYEG complex. This work and our recent study^9^ marks the first time that comparative HDX-MS of membrane proteins was employed to systematically investigate the workings of a multi-subunit protein translocation machinery.

To investigate the conformational dynamics of SecA, we prepared samples of purified SecA alone, saturated with an excess of purified SecYEG. Next we used HDX-MS to interrogate the conformational dynamics of the complex. In each case, the experiments were run in the presence of ADP or a non-hydrolysable analogue of ATP (AMPPNP), and only the data for the saturated component (SecA) was used.

Optimised procedures for HDX-MS were carried out as described previously^9,17^. Following deuteration of the protein, the HDX reaction was quenched at specific time-points (0.25, 1, 5, and 30 mins) by lowering the pH and temperature; the peptides were then digested with pepsin, separated and identified by ultra-performance liquid chromatography (UPLC) and MS (see **Methods**). The identified peptides were then subjected to analysis by MS yielding the mass uptake profiles for each condition (**Figure 1c**). The peptide coverage of the whole complex (ranging from 82 % to 95%) was remarkably high (**Supplementary Figure 1**), enabling a thorough analysis of SecA. The comparison of the AMPPNP *versus* ADP datasets produced a differential HDX-MS fingerprint (ΔHDX) that allowed direct comparisons between the results obtained for distinct states (AMPPNP - ADP; **Figure 1c**). To interpret the data, we utilised the Woods plot prepared using our in-house developed software (Deuteros^25^).

We began by investigating the influence of nucleotides on the conformational dynamics of SecA. We carried out ΔHDX experiments allowing us to track structural changes between AMPPNP and ADP bound states (SecA^AMPPNP^ – SecA^ADP^). Surprisingly, there was very little effect on the conformational dynamics of the isolated SecA (**Figure 2a, b** and **Supplementary Figure 2a**). As expected, the greatest difference in deuterium uptake between the two states was observed at and around the ATP-binding sites (**Figure 2a, b**). This highlights the fact that SecA is only activated when it is associated with SecYEG: unsurprisingly, studying SecA by itself is therefore unlikely to yield information pertinent to transport.

**Figure 2:**
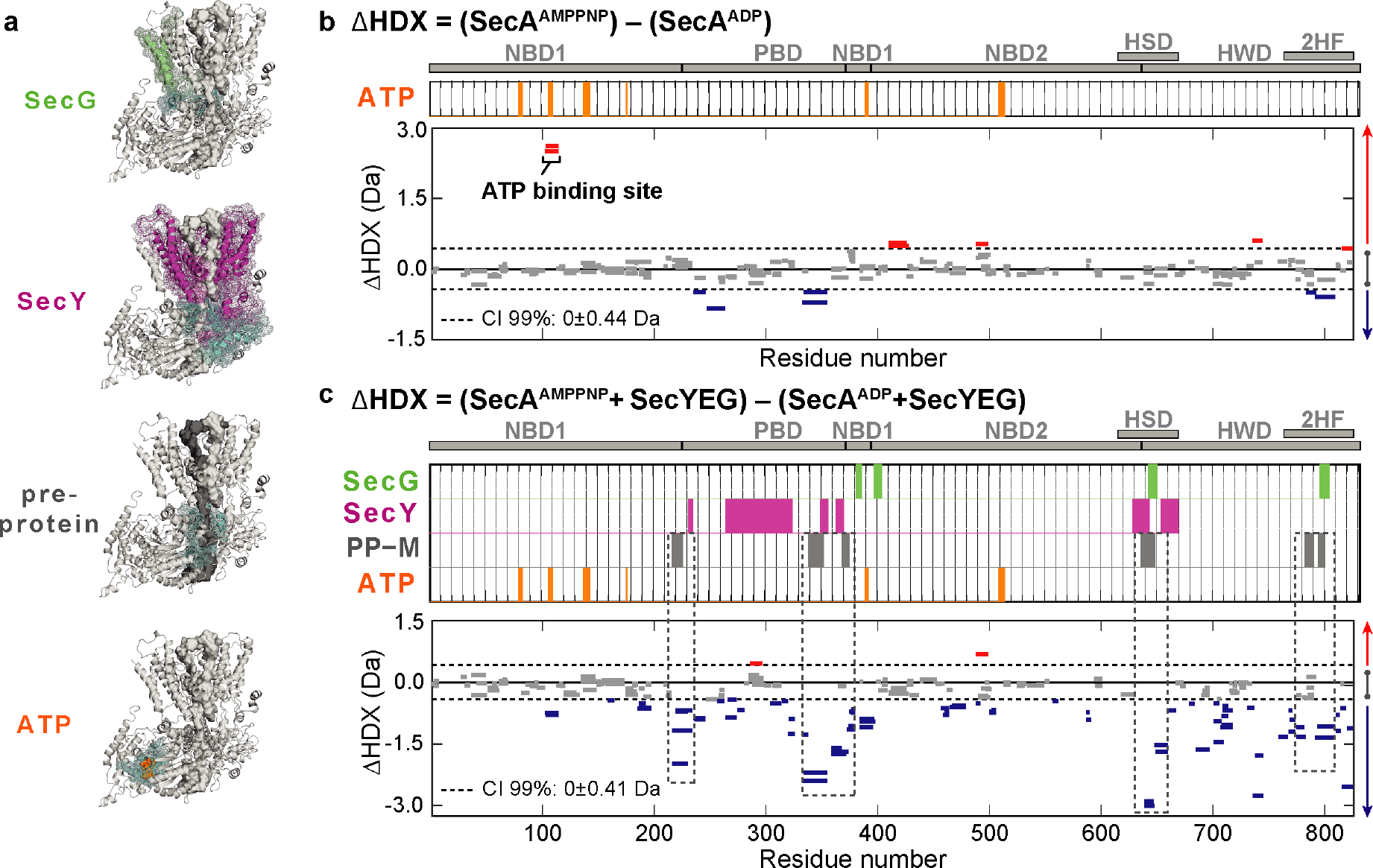
Impact of nucleotides on the conformational dynamics of SecA. **(a)** Structures of the complex highlighting the contact sites between SecA (turquoise) and SecG, SecY, pre-protein and ATP. Significant sum differences in relative deuterium uptake (ΔHDX = AMPPNP-ADP) of (**b**) SecA without SecYEG, and **(c)** of SecA in the presence of excess SecYEG at 30 minutes of deuteration. Highlighted regions represent contacts with SecG (green), SecY (pink), pre-protein (mature domain, grey, PP-M) and ATP (orange).

Therefore, to explore the mechanism of protein transport we carried out equivalent comparative experiments in the presence of an excess of SecYEG to saturate SecA (SecA^AMPPNP^SecYEG - SecA^ADP^SecYEG) (**Figure 2a, c** and **Supplementary Figure 2b**). The results are consistent with SecYEG having a profound effect on SecA: it induces a large ATP-dependent stabilisation across almost the entirety of the protein (**Figure 2c**; negative ΔHDX, blue bars). For mechanistic interpretation, the data were analysed in the context of a model of the SecA-SecYEG bound to a short pre-protein ^26^(**Figure 2a**). Most striking was the correspondence of the stabilised regions with sites in SecA that directly contact a translocating pre-protein (**Figure 2a, c**; dashed boxes –PP-M, gray bars). Such stabilisation in SecA suggests an ATP-driven conformational closure around the mature segment of the pre-protein. Conversely, of course, the highlighted regions identified in SecA would be destabilised in the ADP-bound state.

Next, in order to explore the nucleotide dependent conformational dynamics of the complex we compared the behaviour of SecA to the associated channel complex SecYEG. Previously, we conducted ΔHDX analysis of SecYEGA, as above, but instead saturated SecYEG with SecA^9^. In that instance we were only interested in select peptides lining the cytosolic half of the protein-channel. Here, we focus on the 30 min deuterium exchange (not previously shown; **Figure 3**). When global protein changes are taken into account, ΔHDX (SecA^AMPPNP^SecYEG - SecA^ADP^SecYEG) highlights the long-range nucleotide dependent impact of SecA on the dynamics of SecYEG.

**Figure 3:**
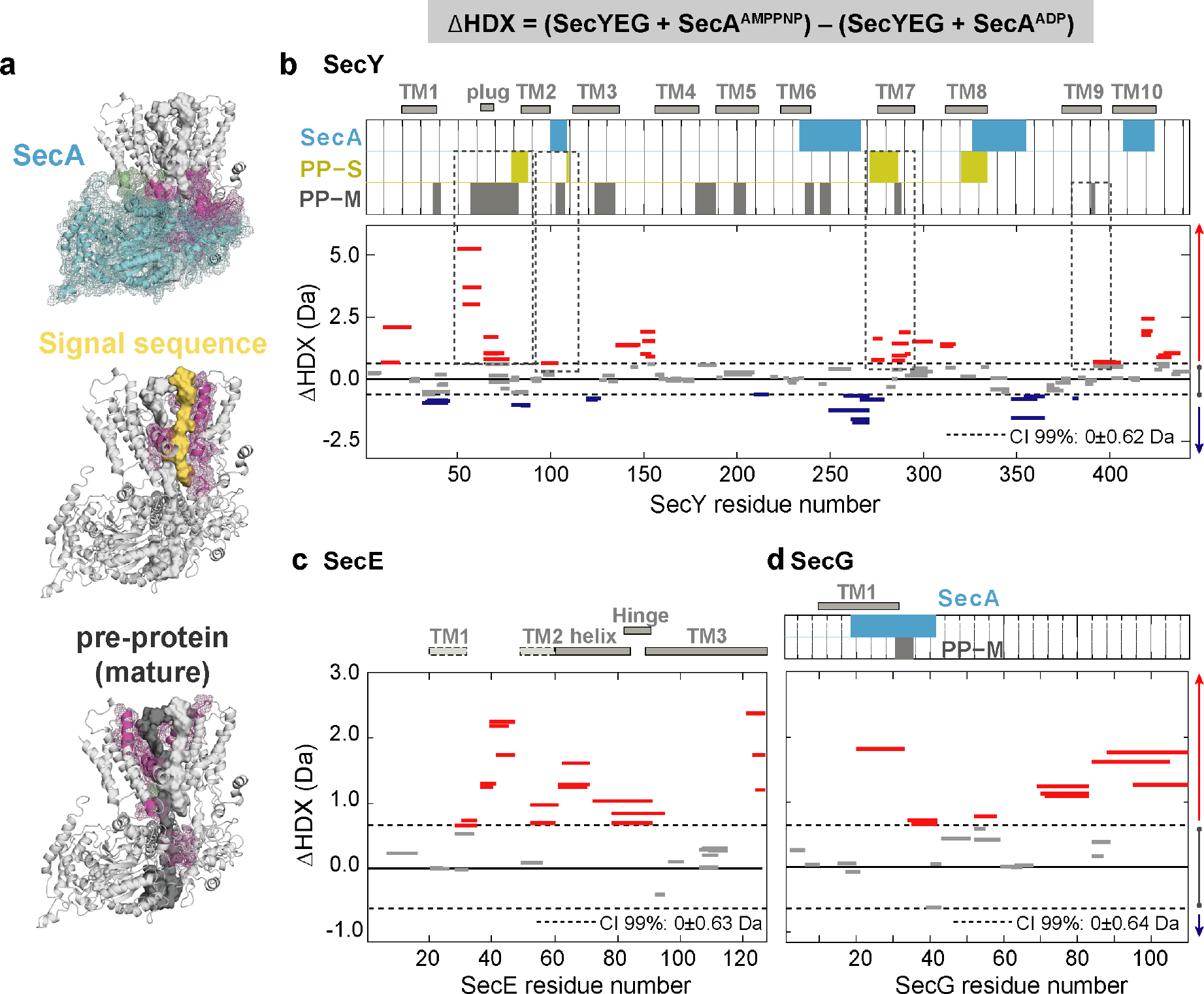
Impact of nucleotides on the conformational dynamics of SecY, SecE and SecG. **(a)** Structures of the SecA-SecYEG complex with regions of SecY (pink) and SecG (green) that contact SecA, signal sequence and pre-protein highlighted. (**b-d**) Significant sum differences in relative deuterium uptake (ΔHDX = AMPPNP-ADP) of **(b)** SecY, **(c)** SecE and **(d)** SecG in the presence of excess SecA at 30 minutes of deuteration. Regions interacting are highlighted: SecA contact (turquoise), pre-protein binding (mature domain, grey, PP-M) and pre-protein binding (signal sequence, yellow, PP-S). Confidence intervals (CI; 99%) are shown as grey dotted lines. Red and blue bars indicate structural stabilisation (positive ΔHDX) and destabilisation (negative ΔHDX) of peptides, respectively. Grey bars indicate peptides with insignificant ΔHDX.

Interestingly, we found significant destabilisation of SecY upon AMPPNP binding (**Figure 3a, b**, and **Supplementary Figure 3a**). Closer inspection of the identified peptides reveals that two prominently destabilised regions in the SecY protein would be in direct contact with the pre-protein during protein translocation (**Figure 3b**; dashed boxes –PP-M gray and PP-S yellow bars). They are the plug, which in the absence of the pre-protein maintains the closed state of the channel^27^, and the highly conserved trans-membrane helix 7. Multiple regions within the SecE and SecG subunits were also significantly destabilised in the presence of AMPPNP (**Figure 3a, c-d** and **Supplementary Figure 3b-c**). Again, conversely those identified regions would be stabilised after hydrolysis to ADP. These observations are consistent with an ATP-driven opening of the SecY channel, and closure after hydrolysis ^7,8^.

The ΔHDX analysis describes contrasting behaviour between the membrane channel and cytoplasmic motor components of the bacterial translocon, during the ATP binding and hydrolysis cycle. Notably, there is an ATP-dependent closure around the translocating polypeptide by SecA and opening of the channel through SecYEG (**Figure 4a**); reversed in the ADP bound state following ATP hydrolysis (**Figure 4b**). To further understand this effect with respect to local structural rearrangements in SecA, we re-analysed previously-run all-atom molecular dynamics (MD) simulations of the SecA-SecYE complex engaged with a stretch of pre-protein^9,12^. We probed the size of two distinct pores in the SecA pre-protein channel through which the pre-protein passes (see **Methods** for details). Note that both pores contain residues shown by HDX-MS to be protected in the ATP-bound state (**Figure 2** and **Methods**).

**Figure 4:**
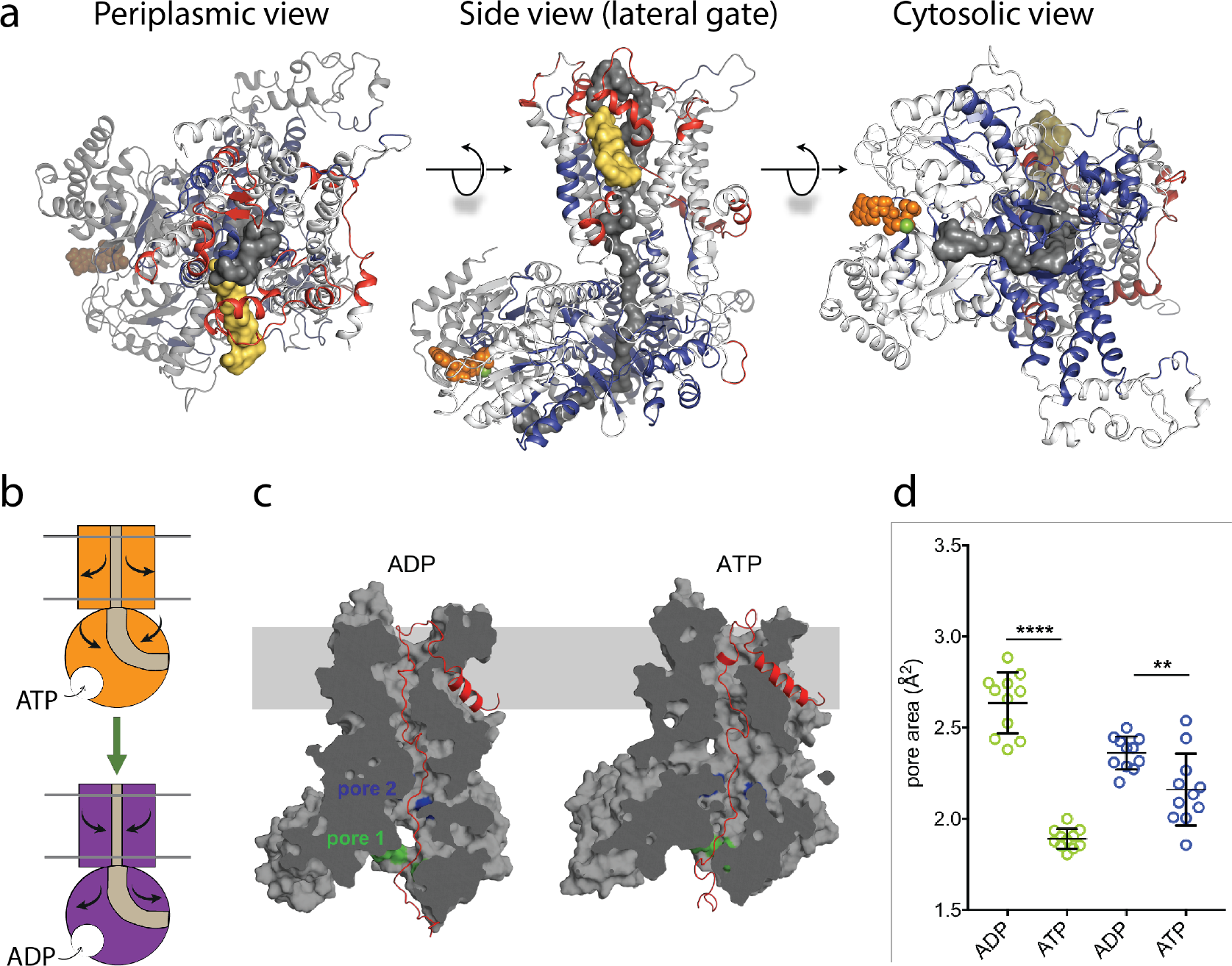
ATP-regulation of channel size. **(a)** Periplasmic, side and cytosolic views of the SecA-SecYEG-preprotein complex model, with ΔHDX mapped on. The translocon complex is coloured light grey, with H-bonding destabilised and stabilised regions in red and blue, respectively. The pre-protein is shown in yellow (signal sequence) and dark gray (mature). ATP is shown as orange spheres. (**b**) Schematic summarising the primary outcomes from the data here. (**c**) Snapshots of SecA-SecYE with an engaged pre-protein after 1 µs all-atom MD simulation^9^. SecA-SecYE is shown as gray surface and has been slabbed to show the pre-protein channel through both SecA and SecY. The pre-protein is shown as red cartoon, and was absent for the cavity size analysis in panel d. The positions of the conserved SecA pores have been highlighted. Visual analysis suggests that pore 1 (green) is more constricted in the ATP state (**d**) Quantification of the pore size, using snapshots every 25 ns from 750—1000 ns. Both pores are tighter in ATP bound state (pore 1 means are 1.9 and 2.6 Å^2^, p < 0.0001: pore 2 means are 2.4 and 2.1 Å^2^, p = 0.0061).

From the simulation data, it is clear that both SecA pores, particularly pore 1 at the cytoplasmic entrance of SecA, are more constricted in the ATP-bound state (**Figure 4c, d**). This is consistent with the HDX data, and suggests that ATP-binding is causing SecA to close around the engaged pre-protein.

Overall, our results reveal wide-ranging differences in the structure of the active Sec complex (SecA-SecYEG) depending on which nucleotide is bound. These differences can broadly be divided into two groups: those affecting the membrane channel (SecY, SecE and SecG), which become deprotected in the presence of ATP, and those affecting SecA, which become protected. A more detailed analysis reveals that the most dramatic changes affect regions that would directly contact a translocating pre-protein. The SecYEG results are consistent with the previously proposed Brownian ratchet mechanism of pre-protein transport, while the observation that ATP causes SecA to constrict around the pre-protein, potentially explaining why pre-proteins do not backslide in the presence of a non-hydrolysable ATP analogue^11,12^. The experiments presented here demonstrate the power of combining HDX-MS – which reveals subtle conformational changes – with the results of biochemistry and molecular simulations to understand the conformational mechanism of multicomponent membrane protein complexes, difficult to study by other approaches.

## Acknowledgment

This work was supported by the Wellcome Trust (109854/Z/15/Z) and a King’s Health Partners R&D Challenge Fund through the MRC, Confidence in Concept (MC_PC_15031) to A.P. This work was funded the BBSRC (BB/N015126/1 to IC; BB/M003604/1 to IC and RAC; BB/I008675/1 to IC and WJA). Extended simulations were run on the ARCHER UK National Supercomputing Service (http://www.archer.ac.uk), provided by HECBioSim, the UK High End Computing Consortium for Biomolecular Simulation (hecbiosim.ac.uk), supported by the EPSRC. EP is the recipient of an Imperial College London Institute of Chemical Biology EPSRC CDT studentship.

