## Supplementary Methods and Figures for "HDX-MS reveals nucleotide-regulated, anti-correlated opening and closure of SecA and SecY channels of the bacterial translocon"

### Supplementary materials and methods

#### Protein preparation

SecYEG and SecA were overproduced and purified as described previously[28].

#### Hydrogen deuterium exchange mass spectrometry (HDX-MS)

The HDX-MS experiments outlined here were carried out using a Synapt G2Si HDMS coupled to an Acquity UPLC M-Class system with HDX and automation (Waters Corporation, Manchester, UK).

To form the SecA-SecYEG complex, SecYEG and SecA were mixed and incubated for 10 min on ice. For experiments investigating SecA (**Figure 2**), SecA (10 µM) was saturated by SecYEG (15 µM). For experiments investigating SecYEG (**Figure 3**), SecA (15 µM) was added in excess of SecYEG (10 µM), as described previously[9]. 5 µl of SecA or the SecA-SecYEG complex was diluted into 95 ml of deuteration buffer (20 mM Tris pH 8, 2 mM $MgCl_2$, 50 mM KCl, and 0.02 % (w/v) dodecyl-maltoside DDM in $D_2O$) or with equilibration buffer (20 mM Tris pH 8, 2 mM $MgCl_2$, 50 mM KCl, and 0.02 % (w/v) DDM in $H_2O$). Deuteration was carried out at 25°C. For experiments analysing the Sec complex in the presence of the nucleotides, 1 mM of either the non-hydrolysable AMPPNP or ADP was added to the protein mixture and to equilibrium or deuteration buffer.

The proteins were labelled by incubation in deuteration buffer for 0.25, 1, 5, and 30 minutes to capture short, medium and long exchange times. Deuteration was quenched with 100 µL of quench buffer (0.7 % (v/v) formic acid and 0.1 % (w/v) DDM) at 1 °C and a pH of 2.4. Protein digestion to peptides was performed at 25°C using an Enzymate online digestion column

(Waters) in 0.1 % (v/v) formic acid at a flow rate of 200 mL/min. Between injection of samples, the pepsin column was washed with cleaning solution (0.8 % (v/v) formic acid, 1.5 M Gu-HCl, and 4 % (v/v) MeOH) recommended by the manufacturer. To reduce peptide carry-over, a blank run was performed between sample runs.

Peptides were trapped using an Acquity BEH C18 1.7 μM VANGUARD pre-column for 3 mins at a flow rate of 200 μL/min in a buffer A of 0.1 % (v/v) formic acid ~ pH 2.5. Peptides were eluted into an Acquity UPLC BEH C18 1.7 μM 1.0 x 100 mm analytical column with a linear gradient of 8–40 % (v/v) gradient of acetonitrile with 0.1 % (v/v) formic acid with a flow rate of 40 μL/min. Peptides were then ionized by positive electrospray into a Synapt G2-Si mass spectrometer (Waters). A 20-30 V trap collision energy ramp was utilised to capture the $MS^E$ data. Electrospray ionization source was operated in a positive ion mode and ion mobility was enabled for all experiments. Leucine Enkephalin was used as a lock mass for mass accuracy correction and iodide was used for mass spectrometry calibration. All deuterium time points were performed in triplicate.

**HDX data evaluation and statistical analysis**

All experiments, including deuterated time points and reference sample controls, were repeated in triplicate. $MS^E$ data from reference sample controls the complexes were used by the Waters ProteinLynx Global Server 2.5.1 (PLGS) and filtered using DynamX (v. 3.0) to provide sequence identification. The following parameters were used to filter the quality of the peptides: minimum and maximum peptide sequence length of 4 and 25, respectively, minimum intensity of 1000, minimum MS/MS products of 2, minimum products per amino acid of 0.2, and a maximum MH + error threshold of 5 ppm. All spectra generated from the peptides were examined and only peptides with a high quality spectra and a high signal to noise ratios were used for data analysis. Woods plots and confidence intervals were generated using the in-house Deuteros software[25].

***In silico* analyses of SecA pore dynamics**

Analyses were based on all-atom molecular dynamics simulations of a complex comprising *Bacillus subtilis* SecA, *Geobacillus thermodenitrificans* SecYE and a 76-stretch of pre-protein, built from PDB 5EUL[6]. The simulations were run in an ATP or ADP-bound state, with full details of their set up described previously[9]. For the analyses here, only the atoms corresponding to SecA and SecYE were kept (i.e. the solvent, membrane and pre-protein were removed). Snapshots were taken every 25 ns over a range of 750 to 1000 ns.

Two highly conserved pre-protein pores in SecA were identified: pore 1 on the cytoplasmic surface of SecA (**Figure 4c**: green) and pore 2 close to the SecY binding interface (**Figure 4**c: blue). Pore 1 consists of residues Ile 222, Ser 224, Gly 326, Arg 327, Arg 328 and Ser 340 (in *B. subtilis* numbering; equivalent to *E. coli* residues Ile 224, Ser 226, Gly 346, Arg 347, Arg 348 and Ser 350). Pore 2 consists of Gln 595, Tyr 599 and Gln 736 on SecA (in *B. subtilis* numbering, *E. coli* equivalent are Gln 644, Tyr 648 and Gln 787), and Ile 243, Tyr 245 and Ala 246 on the functionally-important[29] C4 loop of SecY (*G. thermodentrifinicans* numbering, approximately equivalent to Val 246, Tyr 248 and Ala 249 on *E. coli* SecY). Note that residues Gly 346, Arg 347, Arg 348 and Ser 350 in pore 1 and Gln 787 and Ala 249 of pore 2 (all in *E. coli* numbering) are shown to be deprotected in the HDX data (**see Data Availability).**

To quantify the pore size in the different snapshots, cavity cross-sectional area analyses were run using HOLE [30]. The algorithm was set to start from a position in either pore 1 (between the α-carbons of residues Ile 222 and Ser 340) or pore 2 (between the α-carbons of residues Tyr

599 and Ile 243). To account for flexibility in the pore position, a region of ~0.9 nm on either side of the starting point was selected for analysis, and the pore defined as the narrowest point in this region. The pore size was quantified based on the average over a 0.35 nm window, to reduce artefacts arising from thermal fluctuations.

Images were made using the PyMOL Molecular Graphics System version 2.1.1, Schrödinger, LLC, and data were plotted and analysed in Prism version 7, GraphPad Software.

**Data availability**

All the deuterium uptake plots and datasets from the HDX-MS experiments described in the study are available on figshare data repository.

SecYEG data can be accessed using the following link: xxxxx

SecA data can be accessed using the following link: xxx

SecA-SecYEG (saturated SecA) data can be accessed using the following link: xxxx

SecYEG-SecA (saturated SecYEG from Corey *et al.*, eLife 2019;8:e41803 doi: 10.7554/eLife.41803) data can be accessed using the following link: xxxx

The mass spectrometry proteomics data have been deposited to the ProteomeXchange Consortium via the PRIDE[31] partner repository with the dataset identifier xxxxx.

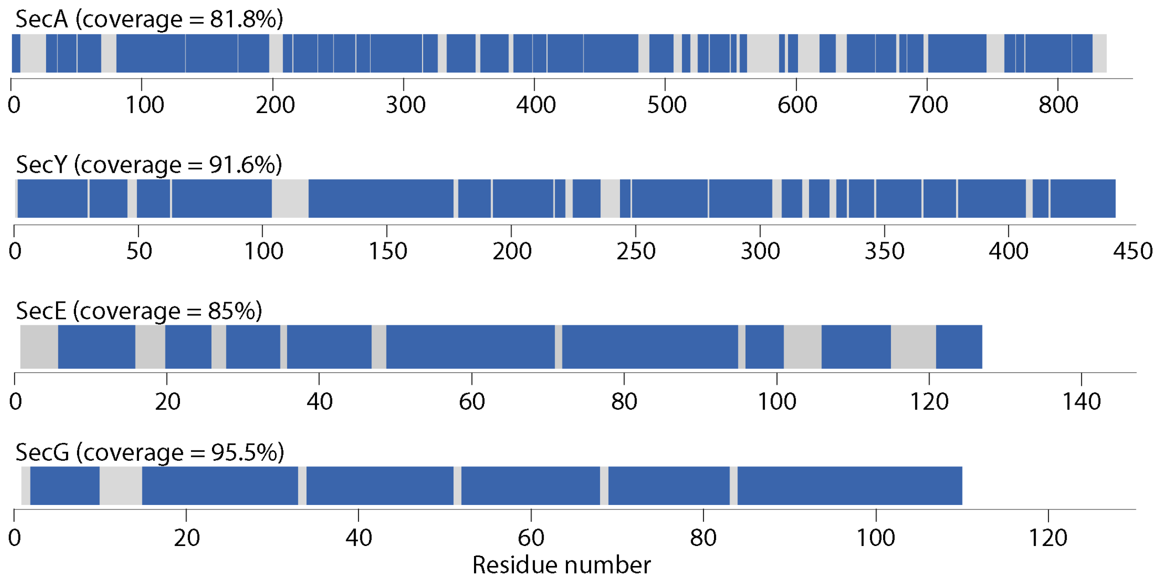


**Supplementary Figure 1.** Linear protein coverage maps for SecA, SecY, SecE and SecG. Maps were generated using Deuteros[25].

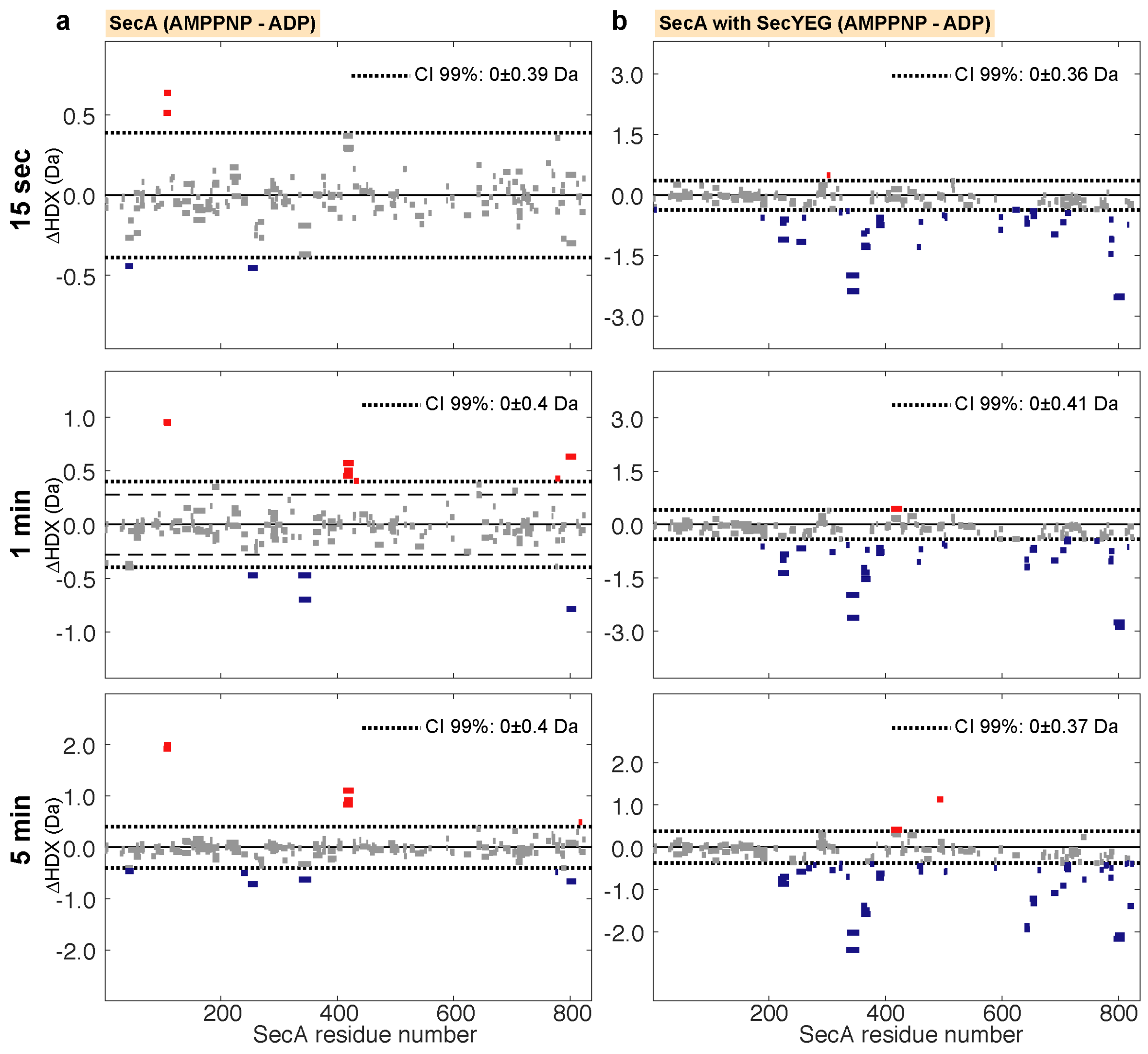


**Supplementary Figure 2.** Woods plots comparing the relative deuterium uptake (AMPPNP-ADP) of (a) SecA alone and (b) SecA in complex (and saturated) with SecYEG after incubation at 25°C in deuterated solvent for 15 seconds, 1, or 5 minutes. The dotted lines represent the 99% confidence interval, which indicates the level of difference of deuterium uptake between the two compared states is statistically significant. Bars represent individual peptides. Bar length corresponds to peptide size. Red and blue coloured bars indicate statistically significant deprotected or protected peptides, respectively.

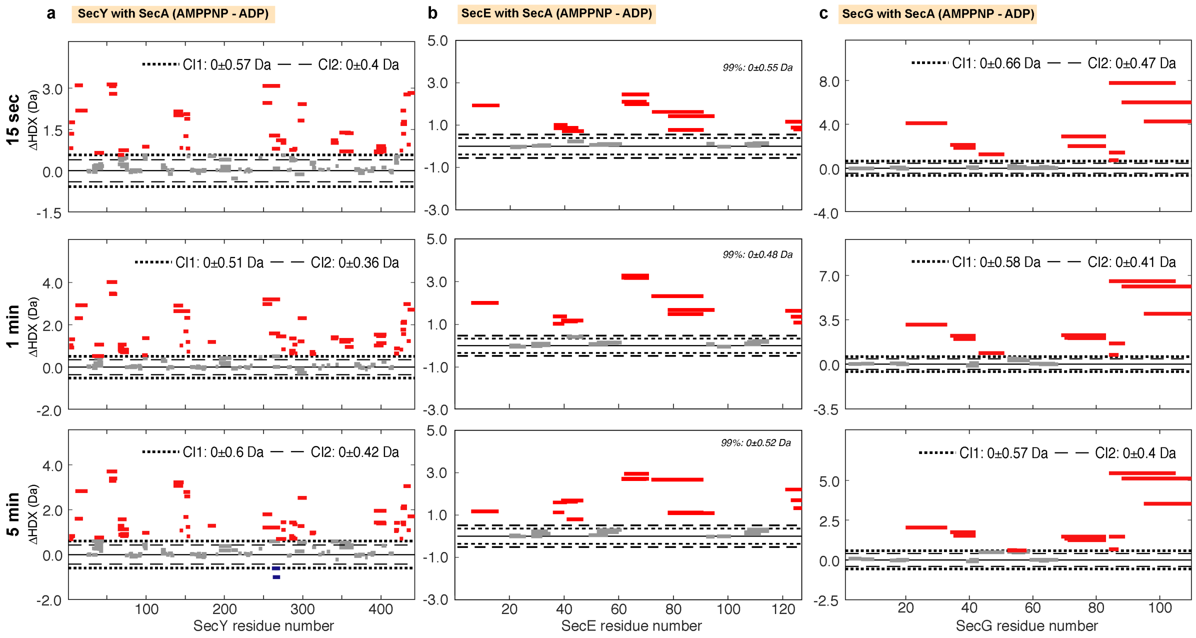


**Supplementary Figure 3.** Woods plots comparing the relative deuterium uptake (AMPPNP-ADP) of (a) SecY (b) SecE and (c) SecG all in complex (and saturated) with SecA after incubation at 25°C in deuterated solvent for 15 seconds, 1, or 5 minutes. The dotted lines represent the 99% confidence interval, which indicates the level of difference of deuterium uptake between the two compared states is statistically significant. Bars represent individual peptides. Bar length corresponds to peptide size. Red and blue coloured bars indicate statistically significant deprotected or protected peptides, respectively.
